# Automatic Whole Body FDG PET/CT Lesion Segmentation using Residual UNet and Adaptive Ensemble

**DOI:** 10.1101/2023.02.06.525233

**Authors:** Gowtham Krishnan Murugesan, Diana McCrumb, Eric Brunner, Jithendra Kumar, Rahul Soni, Vasily Grigorash, Anthony Chang, Jeff VanOss, Stephen Moore

**Affiliations:** BAMF Health, Grand Rapids, Michigan 49503, USA

**Keywords:** CT, Abdominal organ segmentation, Multiple organs, nnUNet

## Abstract

Multimodal Positron Emission Tomography/Computed Tomography (PET/CT) plays a key role in the diagnosis, staging, restaging, treatment response assessment, and radiotherapy planning of malignant tumors. The complementary nature of high-resolution anatomic CT and high sensitivity/specificity molecular PET imaging provides accurate assessment of disease status [14] In oncology, 18-fluorodeoxyglucose (FDG) PET/CT is the most widely used method to identify and analyze metabolically active tumors. In particular, FDG uptake allows for more accurate detection of both nodal and distant forms of metastatic disease. Accurate quantification and staging of tumors is the most important prognostic factor for predicting the survival of patients and for designing personalized patient management plans. [8,3] Analyzing PET/CT quantitatively by experienced medical imaging experts/radiologists is time-consuming and error-prone. Automated quantitative analysis by deep learning algorithms to segment tumor lesions will enable accurate feature extraction, tumor staging, radiotherapy planning, and treatment response assessment. The AutoPET Challenge 2022 provided an opensource platform to develop and benchmark deep learning models for automated PET lesion segmentation by providing large open-source wholebody FDG-PET/CT data. Using the multimodal PET/CT data from 900 subjects with 1014 studies provided by the AutoPET MICCAI 2022 Challenge, we applied fivefold cross-validation on residual UNETs to automatically segment lesions. We then utilized the output from adaptive ensemble highly contributive models as the final segmentation. Our method achieved a 10th ranking with a dice score of 0.5541 in the heldout test dataset (N=150 studies).

## 1 Introduction

Whole-body Positron Emission Tomography/Computed Tomography (PET/CT) is a widely used modality in tumor imaging for evaluating localized tumor burden and detecting symptomatic metastatic lesions early. This non-invasive method allows for the quantification of metabolically active tumors and plays a crucial role in the initial diagnosis, staging, restaging, treatment planning, and recurrence surveillance of various types of cancer. Recent studies have also shown that PET/CT can provide early information on tumor response to therapy, potentially enabling personalized patient management. [2]

PET-based radiotracers are widely used in the diagnosis and management of various types of malignant tumors. Among these radiotracers, 18-fluorodeoxyglucose (18F-FDG) is the most commonly used for oncologic imaging. This is due to the fact that 18F-FDG is based on the increased glucose metabolism in malignant tumors. [12,6] The conventional tracer 18F-FDG has been found to be highly effective in detecting lesions that maintain high glucose metabolism, both in the primary tumor and in metastases. In particular, 18F-FDG has been shown to have a high sensitivity for detecting metastases in solid tumors. [4]

The annotation of lesions is typically performed by expert radiologists to conduct quantitative analysis. However, manual annotation of tumors is a labor-intensive, error-prone, and time-consuming task, especially in whole-body FDG-PET scans. The poor resolution and high statistical noise in PET images, uptake of FDG in several highly metabolic but normal, healthy tissues (e.g., brain and heart) in addition to tumor regions, and a time-dependent blood pool signal, inter-subject uptake variability, and sparse tumor regions in whole-body PET/CT, data acquisition variability, poses further challenges in developing automatic algorithms for tumor segmentation. [7,1] Recent developments in deep learning models that achieve highly accurate PET/CT lesion segmentation in specific regions provide a promising premise to address this issue. A number of recent studies have explored the potential of DL-based automated tumor segmentation from PET or hybrid PET/CT examinations, with a focus on specific disease types or organs such as head and neck cancer, liver, lung, and bone lesions. [13,9,11,10]

The advancements in deep learning models have shown promising results in accurately segmenting PET/CT lesions in specific regions. Several studies have been conducted in the recent past to investigate the potential of DL-based automated tumor segmentation from PET or hybrid PET/CT examinations. These studies have focused on single disease types or organs such as head and neck cancer, liver, lung, and bone lesions. To further advance the state of the art in whole-body PET lesion segmentation, the AutoPET Challenge 2022 provided an open-source platform for developing and benchmarking deep learning models by providing large open-source whole-body FDG-PET/CT data. Our study utilized a 3D residual UNET with five-fold cross-validation on the AutoPET data and applied adaptive ensemble to obtain the final results. Our method achieved top-performing results in the challenge.

## 2 Materials and Methods

### 2.1 Data and Preprocessing

The models were trained using whole-body FDG-PET/CT data from 900 patients, including 1014 studies provided by the AutoPET challenge 2022. A held-out dataset consisting of 200 studies, 100 of which originated from the same hospital as the training database and 100 were drawn from a different hospital with a similar acquisition protocol, was used as a test dataset to assess the robustness and generalizability of the algorithm. In the preprocessing step, the CT data was resampled to the PET resolution and normalized. Two experts annotated the training and test data. A radiologist with 10 years of experience in Hybrid Imaging and experience in machine learning research at the University Hospital Tiibingen annotated all data. A radiologist with 5 years of experience in Hybrid Imaging and experience in machine learning research at the University Hospital of the LMU in Munich annotated all data. We held out 194 studies from unique patients as an internal held-out dataset and used the remaining 1013 studies for five-fold cross-validation. The residual UNET model was trained on five-fold cross-validation using the training set and uploaded into the challenge portal for testing in docker format.

### 2.2 Model Training Methodology

#### Model Architecture

The nnUNET pipeline has achieved top tier performance in multiple medical imaging segmentation competitions. Analysis of the nnUNET pipeline and model architecture has shown that different variations sometimes perform better than the baseline nnUNET architecture [10] [5]. From this, a standard a variant model using residual connections was proposed for training (see Fig. 2 and 3). The input image size of 64×160×160 with one channel, CT is used as input. Input is resampled down five times by convolution blocks with strides of 2. On the decoder side, skip connections are used to concatenate the corresponding encoder layers to preserve spatial information. Instance normalization and leaky ReLU activation in the network layers was used. This architecture initially used 32 feature maps, which then doubled for each down sampling operation in the encoder (up to 1024 feature maps) and then halved for each transposed convolution in the de-coder. The end of the decoder has the same spatial size as the input, followed by a 1×1×1 convolution into 1 channel and a SoftMax function. Models are trained for five folds with loss function of Dice Sorensen Coefficient (DSC) in combination with weighted cross entropy loss were trained. To prevent overfitting augmentation techniques such as random rotations, random scaling, random elastic deformations, gamma correction augmentation, mirroring and elastic de-formation, were adopted. Each of the five models were trained for 1000 epochs with batch size of eight using SGD optimizer and learning rate of 0.01. Dice Similarity Coefficient (DSC), and normalized surface dice (NSD), will be used to assess different aspects of the performance of the segmentation methods.

**Fig. 1.**
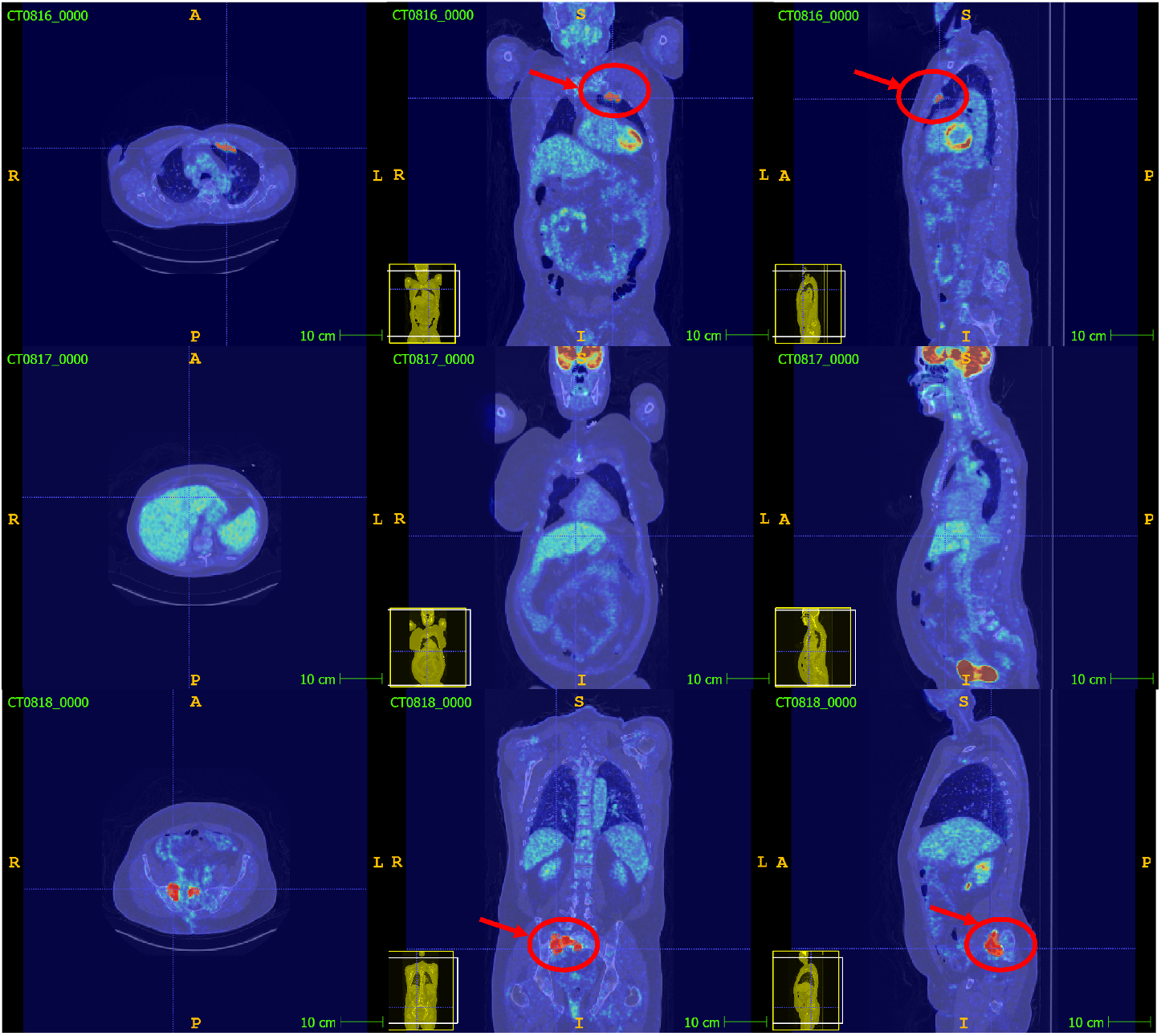
Representative Whole Body FDG PET/CT scans provided by AutoPET challenge with annotations

**Fig. 2.**
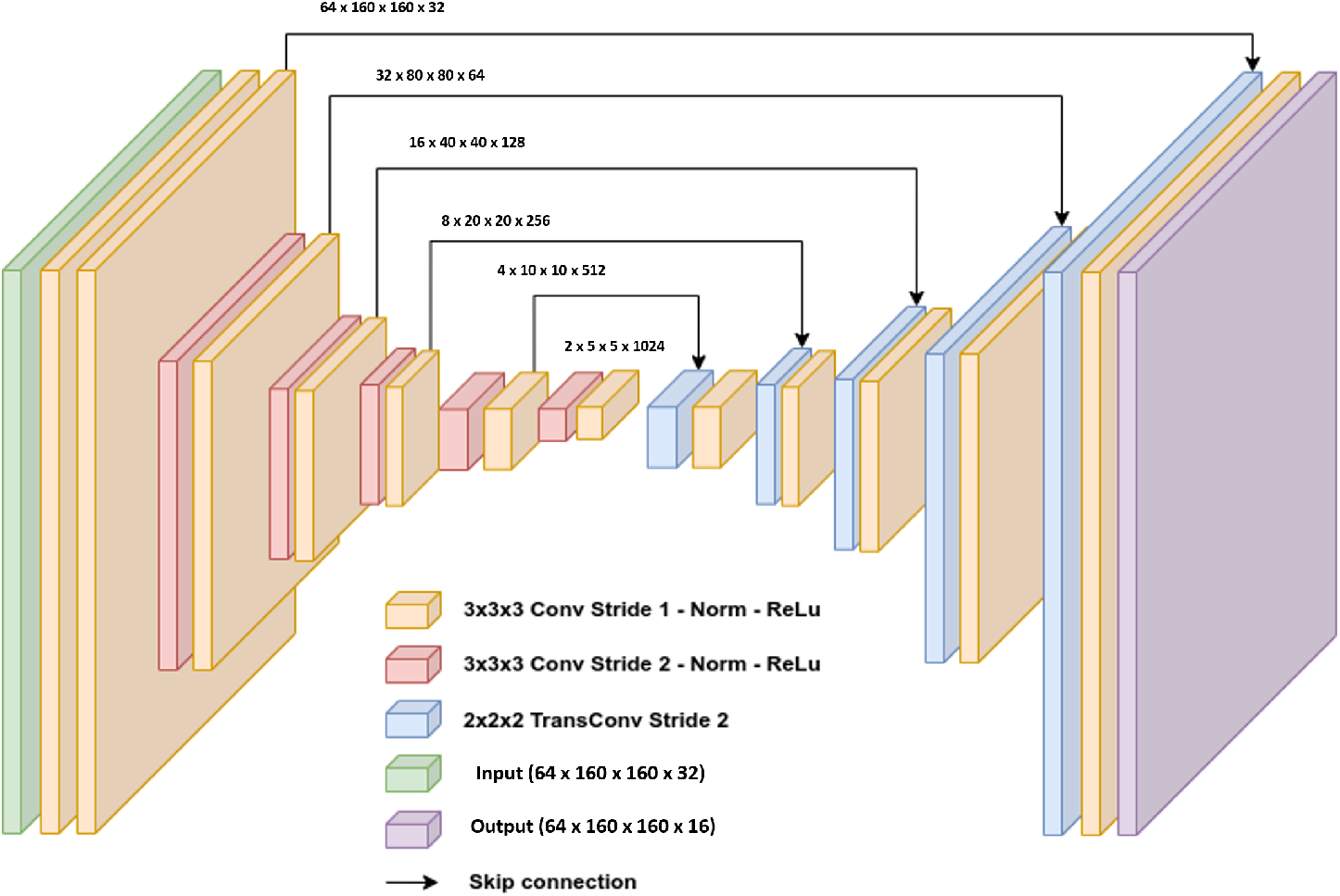
The layers of the UNET architecture used. The input is a volume of 64×160×160 with one channels, CT. Input is resampled down five times by convolution blocks with strides of 2. On the decoder side, skip connections are used to concatenate the corresponding encoder layers to preserve spatial information.

**Fig. 3.**
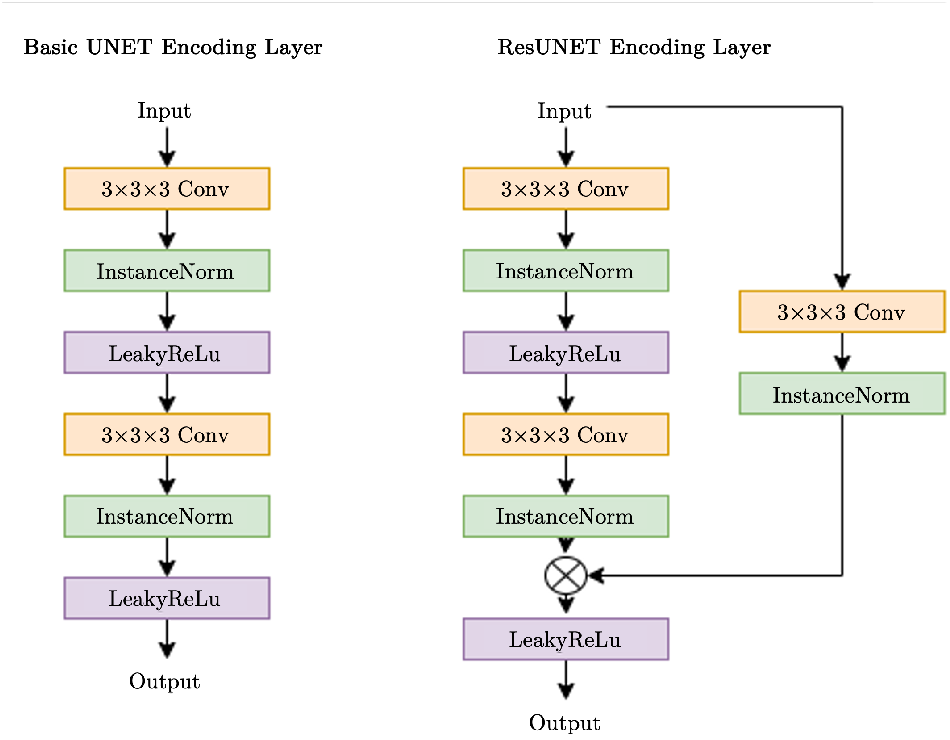
In one instance of our UNET models, each encoding layer is a series of Convolution, normalization, and activation function repeated twice. In another instance, ResUNET, each encoding layer adds a residual path with convolution and normalization.

### 2.3 Results

We trained a five-fold residual UNet model for automatic whole-body lesion segmentation, achieving a robust mean dice score of 0.5541 in the external held-out testing.

### 2.4 Discussion

Our method demonstrated similar performance in both cross-fold validation and unseen held-out data, indicating that it generalized well to multicenter data. The use of adaptive ensemble increased performance by selectively incorporating model outputs with high contribution to the final ensemble. [10] Further improvements in segmentation performance may be achieved through uncertainty-aware segmentation correction.

### 2.5 Conclusion

Our study trained a residual 3D UNet and achieved robust and generalized performance in automatic whole-body FDG-PET/CT lesion segmentation. Our method placed 10th in the MICCAI AutoPET 2022 challenge out of 26 participating teams.

